# Bridging the gap between formal theory and scientific reform practices

**DOI:** 10.1101/2022.12.07.519533

**Authors:** Erkan Buzbas, Berna Devezer

**Affiliations:** Department of Mathematics and Statistical Science, University of Idaho; Department of Business, University of Idaho

**Keywords:** Formal theory, Scientific reform, Reproducibility, Replication, Multi-lab replications, Meta-hypothesis

## Abstract

A core problem that has been addressed in the scientific reform movement so far is the low rates of reproducibility of research results. Mainstream reform literature aimed at increasing reproducibility rates by implementing changes in research practice and scientific policy. At the sidelines of reform, theoreticians have worked on understanding the underlying causes of irreproducibility from ground up. Each approach faces its own challenges. While mainstream focus on swift practical changes has not been buttressed by sound theoretical arguments, theoretical work is slow and initially is only capable of answering questions in idealized setups, removed from real life constraints. In this article, we continue to develop theoretical foundations in understanding non-exact replications and meta-hypothesis tests in multi-lab replication studies, juxtapose these theoretical intuitions with practical reform examples, and expose challenges we face. In our estimation, a major challenge in the next generation of the reform movement is to bridge the gap between theoretical knowledge and practical advancements.

## Introduction

The scientific reform movement that was initiated by a so-called replication crisis has attracted interest of scholars spanning a wide range of disciplines from social and biomedical sciences to humanities and formal sciences. A core problem attacked has been low rates of results reproducibility^1^ in some scientific disciplines—given a result from an original study, the probability of obtaining the same result in replication experiments. First generation approaches predominantly aimed at reforming scientific policy and practice to improve results reproducibility, at times based on hasty diagnoses of problems and normative implementations of quick-and-dirty solutions. This act-first-think-later approach has derailed the investigation of underlying causes of results reproducibility and deprioritized a rigorous theoretical foundation. This has led to the illusion that fundamentally mathematical, complex issues can be solved with relatively simple behavioral fixes. This hesitance to attack a mathematical problem as mathematical seems culturally driven. For example, it should be clear that the true reproducibility rate of a true result in a sequence of exact replication studies with independent and identically distributed random samples can take any value on [0, 1] depending on the elements defining that study. Thus, we should not have a default expectation of high rates of reproducibility or assume that low reproducibility is indicative of false discoveries (Buzbas et al., 2022).

We are statisticians. We expect clear and precise reporting about models and analyses. This clarity and precision of statistical reasoning is largely lacking in the first generation reform literature. We invite the skeptical reader to search through high-profile, large-scale replication studies reporting on results reproducibility and check in how many a formal statement of the statistical model under which inference is made can be identified. We would be in a precarious position to make statistical recommendations without providing a tight statistical reasoning for it. Further, precision is useless without theoretical understanding. As John von Neumann put: “There’s no sense in being precise when you don’t even know what you’re talking about.” Our metascientific research has diverged from the mainstream reform movement as we aim to gain theoretical understanding and offer precision. However, there are outstanding challenges on our side of the world as well.

In this article, we discuss some key statistical issues regarding the reproducibility of scientific results in current large-scale replication studies. Our main point is the importance of distinguishing between exact and non-exact replications and their proper statistical treatment in the analyses. Non-exact replications are determinants of understanding results reproducibility in large scale studies. We evaluate the consequences of this distinction on testing meta-hypotheses in multi-lab-like replication studies using standard statistical theory. And as a next generation challenge, we propose the daunting yet necessary interdisciplinary work of bringing the theory and practice closer together.

## Distinguishing between exact and non-exact replications

We set out to work with an idealized version of the real-life problem. An idealized study comprises background knowledge, an assumed statistical model, and statistical methods to analyze a sample, which is generated under the true model independent of all else (Devezer et al., 2021). A *result* is a function from the space of analysis to the real line. To evaluate the reproducibility rate of a result obtained from a sequence of studies, all replications must randomly sample the same sampling distribution of results. This is guaranteed if a sequence of replication studies are *exact* replications, that is, only the random sample differs through the replications. Statistically, the best way to assess whether a given result from one study is reproduced in its replications is by the relative frequency of that result in all replication studies. A major value of this idealization is advancing understanding of the theory of results reproducibility under exact replications.

In practice, this idealized setup is unrealistic, given uncontrollable sources of variability across replications. No sensible person would argue that they have replicated another study exactly, except perhaps in a textbook probability experiment. When the sampling distribution of the results in replication studies differ from that of the original study or from each other, drawing conclusions about result reproducibility becomes a challenging problem. Leaving aside the (perhaps more important) issue of why the sampling distributions differ from each other, how they differ from each other becomes the primary objective to understand if we are to study results reproducibility properly. Hand-waving at a sequence of non-exact replications as “conceptual replications” and claiming to test the generalizability of an underlying effect won’t do as it is begging the question. Therefore, to develop a broad theory of results reproducibility, we must work with a sequence of not-necessarily-exact replications, and treat the case of exact replications as a special case. Unfortunately, for non-exact replications, we have very limited theory. What we know can be summarized as follows. A sequence of replication studies can be treated as a proper stochastic process. If all the elements of replication studies are exactly equivalent to each other except the sample generated under the true model, then the case of exact replications can be treated as a special case, yielding a single sampling distribution of reproducibility rate. When replications are non-exact, there is no single true reproducibility rate for results from all studies and therefore, no single sampling distribution of reproducibility rate. The mean of the process, then, becomes the target, but it needs to be interpreted carefully (see Buzbas et al., 2022, for more information).

A critical matter is that theoretical conclusions about reproducibility depend on the choice of the *result*, and hence the mode of statistical inference in studying results reproducibility. Harking back to L. J. Savage, we believe that the *result* needs to be “as big as an elephant” to allow for generalizability of conclusions. The least useful mode of statistical inference to study results reproducibility is the null hypothesis significance test under the frequentist approach due to its inflexible interpretation of results probabilistically (e.g., cannot talk about probability of a result unconditional of a hypothesis) and its forced dichotomization of results in hypothesis testing (e.g., reject or fail to reject). These properties obscure the measurable quantitative signal in studies of reproducibility, thereby decreasing the resolution in results which could be used to understand the mechanisms driving irreproducibility. If there exists a problem of result reproducibility in hypothesis testing, it will likely exist under other major modes of inference such as point estimation, interval estimation, model selection, and prediction of future observables.

An efficient way to investigate results reproducibility information theoretically is under a more mathematically refined mode of inference than null hypothesis significance testing. Estimation theory is the best understood mode of statistical inference, equipped with most well-established theorems. It is our safety net. Based on estimation theory, model selection is a challenging but ultimate target in modern statistics (as well as in Devezer et al., 2019).

Now that we have laid out our foundations, let us turn to the theoretical challenges that arise when trying to understand results reproducibility from non-exact replications.

## Evaluating the results from non-exact replications with respect to results reproducibility

We assume that a true data generating mechanism *M*_*T*_ that we would like to fully identify exists^2^. For convenience, we assume a linear model with additive errors *M*_*T*_ := *{Y* |**X**_*T*_ *β*_*T*_ = **X**_*T*_ *β*_*T*_ + *ϵ}*, which accommodates common research studies. Here, *Y* is *n ×* 1 vector of response variables, **X**_*T*_ is *n × p* matrix of predictors (e.g., design matrix in experiments), *β*_*T*_ is *p ×* 1 vector of model parameters, and *ϵ* is *n ×* 1 vector of stochastic errors. It is convenient to assume zero mean, constant variance, and uncorrelated errors, which are collectively known as the ordinary least squares (OLS) assumptions. Often, a full model specification is required for analysis, so it is equipped with normal distribution of errors. Thus, *M*_*T*_ is an exactly determined representation of a natural phenomenon of our interest or in other words, the *target true* data generating mechanism. In practice, we may not know the space on which *M*_*T*_ exists. It is worth mentioning that only if the set of models considered by a researcher includes *M*_*T*_, we can hope to study results reproducibility of true results. More precisely, we should be able to include *M*_*T*_ as the saturated model in our consideration with possible null values for parameters. Here, we assume that we can satisfy this assumption. However, we invite the reader to ponder on how challenging it is to satisfy this assumption in practice.

If a study assumes *M*_*T*_ for its analysis, and there was no intervention to affect the data generated by *M*_*T*_, then the solution 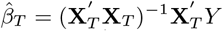 are best linear unbiased estimators. Almost all statistical inferential goals require 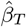 and without loss of generality, here we set it as our *result*.

A crucial question is: If *M*_*T*_ is the true model, how does *M*_*O*_, the assumed model in the original (first) study differ from *M*_*T*_, and how does *M*_*R*_ the assumed model in a subsequent non-exact replication study differ from *M*_*O*_ ? Under different study conditions, we expect the natural phenomenon represented by *M*_*T*_ to be distorted in some way. For example, a controlled study imposes constraints. Two studies can rarely be guaranteed to have identical assumed models that are also identical to *M*_*T*_. This condition requires that all variables that are indeed in *M*_*T*_ to be included and no variable that is not in *M*_*T*_ to be excluded in the assumed models of both studies. We can note the consequences of the violations of this condition on estimates as follows.

We assume that the original study includes only some of the variables that are indeed in *M*_*T*_, and also include some variables that are not part of *M*_*T*_ into the assumed model. The relevant matrix of predictors (e.g., design matrix) which we denote by **X**_*O*_ in the assumed model *M*_*O*_ has now changed. If it is correctly identified, this study operates under the data generating model *M*_*O*_ := *{Y* |**X**_*O*_*β*_*O*_ = **X**_*O*_*β*_*O*_ + *ϵ*}, and the solution for OLS estimates in *M*_*O*_ is now 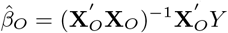. There is no convincing argument to assume that *M*_*O*_ is equivalent to *M*_*T*_. Thus, *β*_*T*_ ≠ *β*_*O*_ implying that 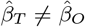 given the same data. Even if *M*_*O*_ is equivalent *M*_*T*_, it is more likely for the replication studies in the sequence to generate data under models different than *M*_*T*_ or *M*_*O*_ via the same process of inclusion and exclusion of variables as described above. Let the assumed model (if identified correctly) in a replication study be *M*_*R*_ := *{Y* |**X**_*R*_*β*_*R*_ = **X**_*R*_*β*_*R*_ + *ϵ}*, and the solution for OLS estimates in *M*_*R*_ is now 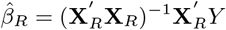. Thus, it would take even a more convincing argument to assume that *M*_*T*_, *M*_*O*_, and *M*_*R*_ are all equivalent to each other.

### A simple example to show drastic effects of non-exact replications

Let us assume that the true model is *M*_*T*_ := *E*(*Y* |*β*_*T*_) = *β*_*T*_, that is a one-parameter model with no predictors. Our interest is to estimate the population mean *β*_*T*_ using the sample *Y*_1_, *Y*_2_, …*Y*_*n*_ independently generated under the assumed model in the original study *M*_*O*_ := *E*(*Y* |*β*_*O*_) = *β*_*O*_. As we have indicated in general *β*_*o*_ ≠ *β*_*T*_. Without loss of generality, let us define a very simple relationship *β*_*O*_ = *β*_*T*_ + *c*, where *c* ≠ 0 is some constant. The best estimator of the population mean is the sample mean and so we use the estimator 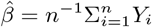. However, we quickly realize that 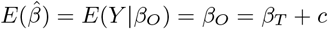. Thus, the sample mean of the data obtained in the study with assumed model *M*_*O*_ is a *biased estimator* for the true population mean of interest *β*_*T*_ under the true model. And it gets worse. We also quickly realize that by weak law of large numbers 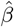 is a consistent estimator of *β*_*O*_ the population mean from which the sample is drawn and *β*_*T*_ = *β*_*O*_ − *c* implies that the probability 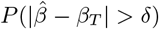 does not converge to 0 for any small *δ >* 0 as the sample size increases. Thus, the sample mean of the data obtained in the study with assumed model *M*_*O*_ is also an inconsistent estimator for the true population mean of interest *β*_*T*_ under the true model. An immediate implication of this argument is on the choice of sample size as the divergent factor between a true model, an original study, and its replications. To assess the reproducibility of a given result, the result must be obtained from studies with equal sample sizes across original and replication studies and identical methods must be applied to arrive to a result. Researchers in practice might be tempted to increase the sample size in a replication (e.g., for higher statistical power as a better standard to meet). However, larger sample sizes do not necessarily imply that true results will be more reproducible. To extend the theoretical example given above, we consider testing the true hypothesis *β*_*T*_ = 0 using intervals 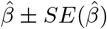, where 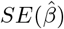 is the standard error of the sampling distribution of 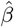. The reproducibility rate, defined as the proportion of the number of true results to the number of total results in a sequence of studies, in this test decreases with larger sample sizes, converging to zero (Figure 1). This leads to a dilemma: Do we strive to conduct very few replication studies but making them as exact as possible at a great expense of resources because we know how to interpret their results theoretically, or do we try and run studies that are non-exact even if we do not quite understand what the implications of this non-exactness may be?

**Figure 1.**
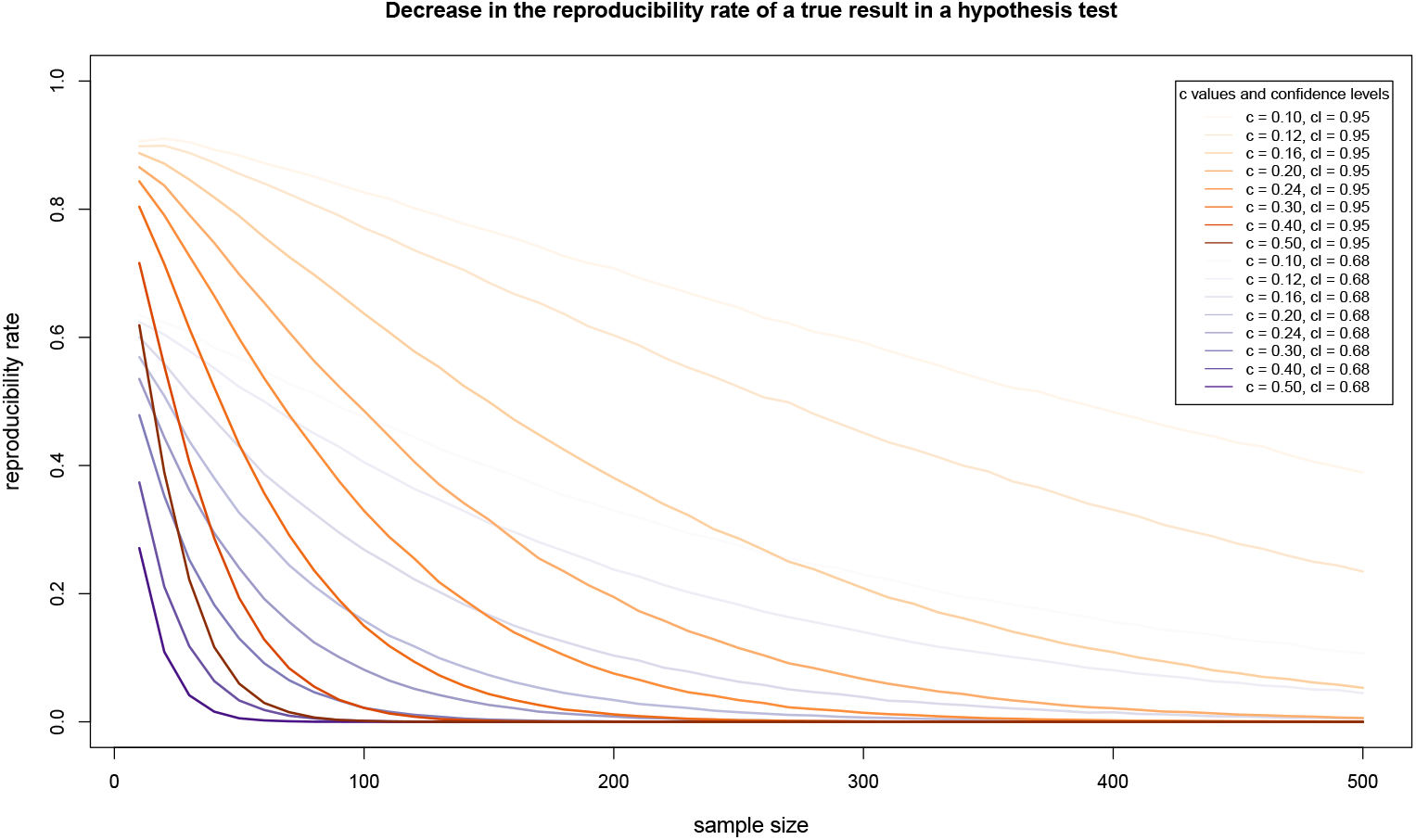
The reproducibility rate of a true result in a hypothesis test decreasing with increasing sample size. The (true) null hypothesis is *β*_*T*_ = 0, where *β*_*O*_ = *β*_*T*_ + *c* and *c >* 0 is a constant.

We conclude that to connect the results *non-exact* replication studies, we need theoretical results on how non-exact they are and on what the exact effect of non-exactness is on the results. Once again, hand-waving at perceived closeness or likeness will introduce further obscurity and make it more difficult to properly interpret the replication results. In Buzbas et al. (2022), we show how easily true or false results can be made more or less reproducible with small perturbations in study design.

Many analyst studies (e.g., Breznau et al., 2022; Hoogeveen et al., 2022; Silberzahn et al., 2018) provide an example of how this type of non-exactness may play out in practice. In these studies, independent research teams are given the same data set and asked to perform analyses to test a given scientific hypothesis. What is striking is that not only inferential procedures but also assumed models vary greatly across analysts, resulting in a range of results not necessarily consistent with each other. Note that these studies use the same data set that samples a given population. In non-exact replication studies, inference is performed from independent data sets often sampling different populations. Also worth noting is that the many analyst results confirm our earlier point about the weakness of focusing on hypothesis tests instead of a more generic modeling framework.

## Studies for testing meta-hypotheses under non-exact replications

As a more advanced example to show how the points we presented might affect the conclusions about results reproducibility in replication studies, we consider the following models for testing meta-hypotheses, for example as in multi-lab studies (e.g., author involvement effect tested in Many Labs 4, Klein et al., 2022). For the *i*th observation, we define *g*(*u*_*i*_, *v*_*i*_) as a function of 1 *× p* vector of predictors such that *u* are variables associated with the meta-hypothesis (e.g., varying on larger experimental units) and *v* are variables associated with the original predictors (e.g., varying on smaller experimental units) of the study (Jones and Nachtsheim, 2009). The matrix of predictors **X**_*T*_* has ith row as *g*(*u*_*i*_, *v*_*i*_). Due to two levels of the study, stochastic error must now conform not only the errors within each replication, but also the errors associated with meta-hypothesis. For the meta-hypothesis we have **Z***η* where **Z** is *n × k* of indicator functions whose *k*^*th*^ element is 1 (e.g., for a level of meta-hypothesis) and others 0, and *η* is *k ×* 1 vector of normally distributed errors with 0 mean and 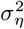 variance. For replications we have *ϵ* is *n ×* 1 normally distributed errors with 0 mean and 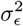 variance for each observation. Assuming additive errors as before, we have **Z***η* + *ϵ* for stochastic errors, and *η*_*i*_ and *ϵ*_*j*_ are assumed to be uncorrelated for all observation pairs (*i, j*). The true model generating the data is *M*_*T*_* := *{Y* |**X**_*T*_* *β*_*T*_* = **X**_*T*_* *β*_*T*_* + **Z***η* + *ϵ}*. Parallel to the argument leading to *M*_*O*_ from *M*_*T*_, the *feasible generalized least squares estimates* in an original study *M*_*O*_* is 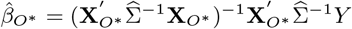, where 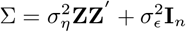 is the covariance matrix. Σ is often unknown. It is estimated by sample variances at the meta-hypothesis and replication variables levels, assuming that the replications are exact.

The estimator 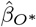 clearly shows why non-exactness of replications will exacerbate errors in estimates in larger models such as those used in testing meta-hypotheses. The reason is that larger models often require nuisance parameters also to be estimated in addition to parameters of interest. Even if we take the best approach to inference, the non-exactness of the replication will be reflected on multiple estimates resulting in undesirable estimators.

For our example, to obtain the estimate of interest 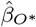, we also need to obtain 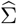. The method of estimation for 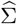 (e.g., restricted maximum likelihood or Bayes) will inevitably use **X**_*O*_* and the properties of 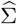 will be affected by non-exactness of replications. This leads to biased estimates. Hence, non-exactness of replications casts more doubt on the inferential results of large models such as meta-hypothesis tests.

Many Labs 4 (Klein et al., 2022) provides a case study. The original hypothesis being tested over replications is the mortality salience effect and the experimental units are individual research participants; the meta-hypothesis being tested is the effect of original authors’ involvement where experimental units are participating labs. The replications are non-exact in many ways from populations being sampled to unequal cell and sample sizes, from variations in in-house protocols to unequal block sizes. All of these factors likely bias effect size estimates in unpredictable ways. Considering the fact that inference is not made under a model that accounts for the randomization restriction and hierarchical experimental units in the actual experimental design, the results become even harder to interpret. The theory tells us what conclusions cannot be supported by the design and the analysis but it does not tell us what conclusions can be justified; that is, until that theory is advanced specifically for such cases.

## Conclusion

We can clearly use mathematical statistics to advance our theoretical understanding of replications and results reproducibility. But whenever theory meets reality, its reach falls short.

We know a lot about the consequences of exact replications yet we also know that exact replications are hard to achieve. In practice, we might think that even if we cannot perform exact replications, controlled lab conditions can approximate the ideal conditions assumed by statistical theory. This is a strong assumption that often gets violated. First, the effect of sampling different populations in replications cannot be remedied by randomization. Second, an approximation is a statement about a quantity approaching to another quantity in a precise mathematical way. It is not some haphazard likeness that we do not know how to define or verify. The mathematical approximation can be measured with precision but the perceived likeness of studies cannot. For example, for the sample mean, we know that the variance of its sampling distribution decreases linearly with the sample size and we can measure the performance of this approximation in an analysis, data point per data point if need be. We would be challenged, however, to show how much a given replication study approximates an original study with respect to a reasonable measure in a statistical sense. The theory for measuring such closeness or likeness does not yet exist. Hence, our theoretical knowledge of exact replications is of little direct use for practice and for a thorough understanding of real life non-exact replications, we lack as rigorous a theory.

Mainstream scientific reform movement has focused on easy-to-implement behavioral changes (e.g., preregistration, code sharing) as well as reforms in scientific policy (e.g., reducing publication bias). As we alluded to earlier, these developments are often based on intuition and not grounded in theoretical understanding. In the same period, progress has been made toward building theoretical foundations of metascience by our own work and others’ (Bak-Coleman et al., 2022; Fanelli, 2022; Fanelli et al., 2022; Smaldino and McElreath, 2016), albeit in the margins of reform. Such theoretically guided approaches, on the other hand, are constrained by the scope of the idealizations involved and may not readily translate to scientific practice. As theoreticians our job does not end with developing rigorous theory. We also strive to reach a “post-rigorous” stage, as mathematician Terence Tao observed, where we have grown comfortable enough with the rigorous foundations of metascience so that we we finally feel “ready to revisit and refine [our] pre-rigorous intuition on the subject, but this time with the intuition solidly buttressed by rigorous theory” (Tao, 2007). To us then the major challenge in the next generation reform is this: How do we bridge the gap between theory and practice?

## Footnotes

1 Also referred to as *replicability* in the literature.

2 Otherwise we would be chasing a moving target. There exist tools to study these cases, but they are out of our scope for the treatment of results reproducibility.

